# Best Practices for Benchmarking Germline Small Variant Calls in Human Genomes

**DOI:** 10.1101/270157

**Authors:** Peter Krusche, Len Trigg, Paul C. Boutros, Christopher E. Mason, Francisco M. De La Vega, Benjamin L. Moore, Mar Gonzalez-Porta, Michael A. Eberle, Zivana Tezak, Samir Labadibi, Rebecca Truty, George Asimenos, Birgit Funke, Mark Fleharty, Brad A. Chapman, Marc Salit, Justin M Zook, and the Global Alliance for Genomics and Health Benchmarking Team

**Author notes:** These authors contributed equally to this work (Marc Salit, Justin M Zook). Contact Information: Justin Zook.

## Abstract

Standardized benchmarking methods and tools are essential to robust accuracy assessment of NGS variant calling. Benchmarking variant calls requires careful attention to definitions of performance metrics, sophisticated comparison approaches, and stratification by variant type and genome context. To address these needs, the Global Alliance for Genomics and Health (GA4GH) Benchmarking Team convened representatives from sequencing technology developers, government agencies, academic bioinformatics researchers, clinical laboratories, and commercial technology and bioinformatics developers for whom benchmarking variant calls is essential to their work. This team addressed challenges in (1) matching variant calls with different representations, (2) defining standard performance metrics, (3) enabling stratification of performance by variant type and genome context, and (4) developing and describing limitations of high-confidence calls and regions that can be used as “truth”. Our methods are publicly available on GitHub (https://github.com/ga4gh/benchmarking-tools) and in a web-based app on precisionFDA, which allow users to compare their variant calls against truth sets and to obtain a standardized report on their variant calling performance. Our methods have been piloted in the precisionFDA variant calling challenges to identify the best-in-class variant calling methods within high-confidence regions. Finally, we recommend a set of best practices for using our tools and critically evaluating the results.

## Introduction

Next generation sequencing (NGS) technologies and analysis methods have rapidly evolved and are increasingly being used in research and clinical settings. The ability to detect DNA variants began in the last third of the 20th century when recombinant DNA technology facilitated the identification and characterization of human genes. Due to the high cost of sequencing technologies, early diagnostic applications were limited to screening patient samples for established pathogenic variants. Clinical heterogeneity and overlapping presentations can complicate accurate diagnosis based on clinical symptoms alone, which often resulted in the need for sequential testing approaches (diagnostic odysseys). While some focused tests are still in use today, large gene panels including tens to hundreds of genes, often accommodating sets of diseases with clinical overlap, are the most common application for NGS today, with exome and genome sequencing rapidly gaining popularity in the research and medical genetics communities.^1,2^ An output of these tests is a list of variant calls and their genotypes, often in variant call format (VCF), and benchmarking these calls is an important part of analytical validation.

Robust, sophisticated, and standardized benchmarking methods are critical to enable development, optimization, and demonstration of performance for sequencing and analysis tools. This is especially important for clinical laboratories developing sequencing-based tests for medical care. Efforts such as the *Genome in a Bottle Consortium* and *Platinum Genomes Project* have developed small variant “truth” sets for several well-characterized human genomes from publicly available cell lines and DNA.^3–6^ A “truth” set was also recently developed from a “synthetic-diploid” mixture of two haploid hyditaform mole cell lines not currently in a public repository.^7^ A framework for benchmarking non-complex small variant calls in the exome was developed previously as a web-based tool GCAT.^8^ However, comparing variant calls from any particular sequencing pipeline to a truth set is not a trivial exercise. First, benchmarking must consider that variants may be represented in multiple ways in the commonly used variant call format (VCF).^9–12^ When comparing VCF files record by record, many of the putative differences are simply different representations of the same variant. Secondly, definitions for performance metrics such as true positive (TP), false positive (FP), and false negative (FN), which are key for the interpretation of the benchmarking results, are not yet standardized. Lastly, due to the complexity of the human genome, performance can vary across variant types and genomic regions, which inevitably increases the number of benchmarking statistics to report.

In the context of performance metrics, two critical performance parameters that are traditionally required for clinical tests are sensitivity (the ability to detect variants that are known to be present or “absence of false negatives”, which we call “recall” in this work) and specificity (the ability to correctly identify the absence of variants or “absence of false positives”, which we replace with “precision” in this work).^13^ The shift from focused genotyping tests to genome sequencing enables the detection of novel sequence variants, which has fundamental implications on how these diagnostic performance parameters need to be determined. Early professional guidelines call for the use of samples with and without known pathogenic variants to determine sensitivity and specificity, which was appropriate when genetic testing interrogated only targeted, previously identified variants. This approach remains valid for sequencing-based testing, but now constitutes an incomplete evaluation, since it does not address the ability to detect novel variants. To predict performance for novel variants, it is important to maximize the number and variety of variants that can be compared to a “gold standard” in order to establish statistical confidence values for different types of variants and genome contexts, which can then be extrapolated to all sequenced bases.^14–17^ While this problem already existed for Sanger sequencing tests, the power and scope of NGS technologies presents a different scale of challenges for fit-for-purpose test validation. Laboratories that performed Sanger sequencing prior to transitioning to NGS were often able to utilize previously analyzed specimens to establish analytical performance of NGS tests; however, this approach is practically limiting, poses severe challenges for other laboratories, and is completely infeasible as test sizes increase from a few genes to the exome or genome. Guidelines were recently published for validating clinical bioinformatics assays.^18^ These guidelines highlight the utility of reference materials for benchmarking variant calls, as well as the importance of stratifying performance by variant type and genome context.

To address the needs for using reference materials to benchmark variant calls in a standardized, robust manner, we present the work of the Global Alliance for Genomics and Health (GA4GH) Benchmarking Team. This team, open to all interested parties, includes broad stakeholder representation from research institutes and academia, sequencing technology companies, government agencies, and clinical laboratories, with the common goal of driving towards the standardization of variant calling benchmarking. In particular, we describe the available reference materials and tools to benchmark variant calls, and provide best practices for using these resources and interpreting benchmarking results.

## Results

Our goal was to standardize the variant benchmarking process such that (1) the methods used to compare callsets assess the accuracy of the variant and genotype calls independent of different representations of the same variant, (2) primary performance metrics are represented in the most commonly used binary classification form (i.e., TP, FP, FN, and statistics derived from these), (3) calculation of performance metrics is standardized such that they can be compared more easily across methods, and (4) performance metrics can be stratified by variant type and genome context.

We discuss the technical challenges presented by comparing VCF files accurately and describe our solution to implement such comparisons. We focus on the use case where we have a call set that can be used as “truth” (e.g., Genome in a Bottle or Platinum Genomes) and would like to benchmark a single-sample query VCF against this dataset. The inputs to this comparison are a *truth callset* (in VCF format), and a set of *confident regions* (in BED format) for the truth set. The confident regions indicate the locations of the genome where, when comparing to the truth callset, variants that do not match the truth callset should be false positives and variants missed in the truth callset should be false negatives. Furthermore, our inputs include a *query callset* in (g)VCF format, a reference FASTA file and optionally stratification regions to break out variant calling performance in particular regions of the genome or to restrict comparisons to a genomic subset (e.g. exons / regions captured by targeted sequencing). For more details see SI A. We developed a framework for standardized benchmarking of variant calls (Fig. 1), which addresses the challenges discussed in detail in the following sections.

**Figure 1:**
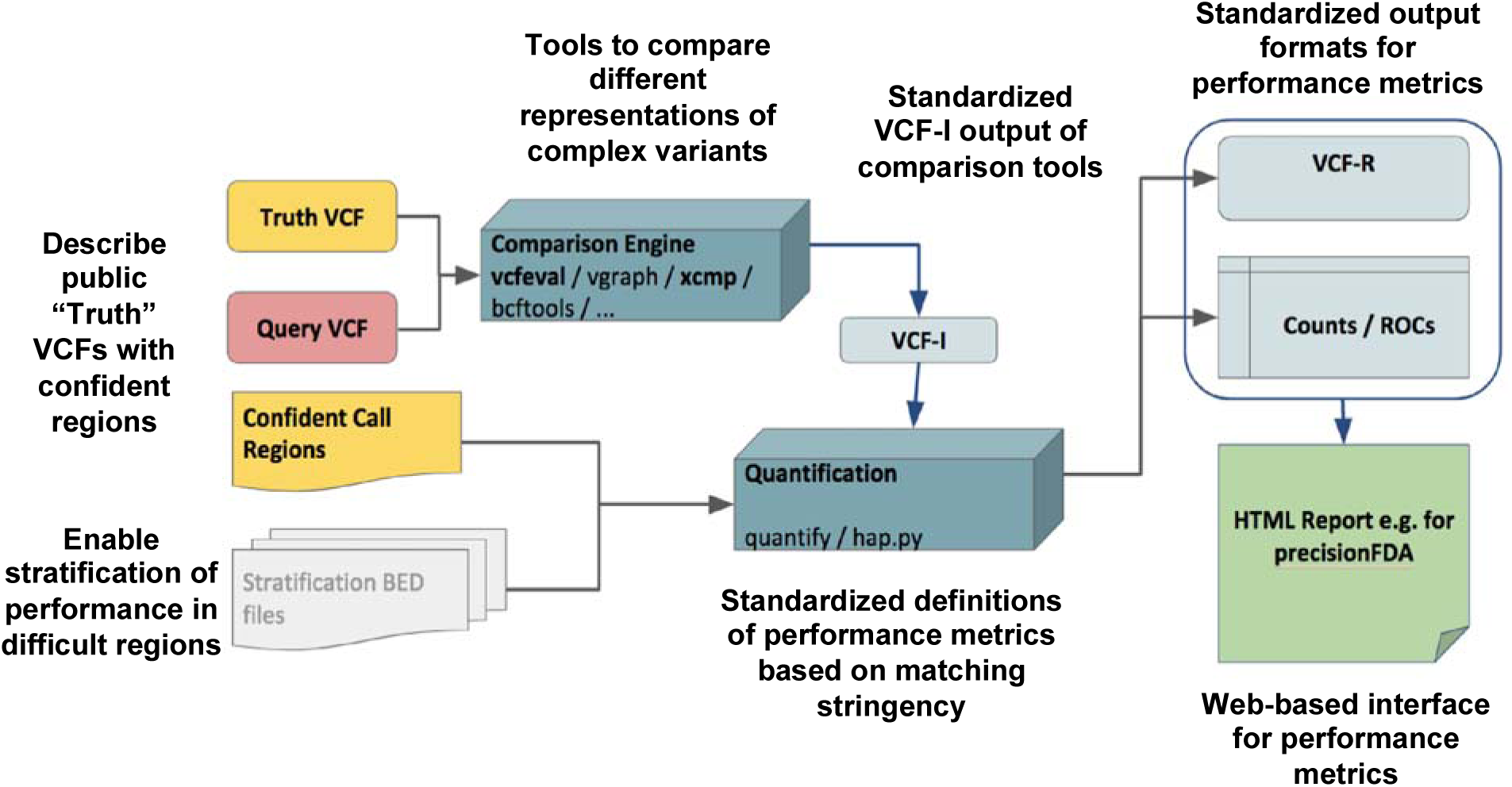
The GA4GH Benchmarking Team’s reference implementation of a comparison framework, annotated with free-floating text describing the team’s innovations. The framework takes in a Truth VCF, Query VCF, confident call regions for the Truth and/or Query, and optionally BED files to stratify performance by genome context. A standardized intermediate output (VCF-I) from the comparison engines allows them to be interchanged and for TP, FP, and FN to be quantified in a standard way.

## Variant representation

The primary challenge with comparing two VCF files is handling complex variant representations correctly. In a VCF file, we describe two haplotype sequences by means of REF-ALT pairs and genotypes. These variant calls do not always uniquely represent the same haplotype sequences. Alignments are not always unique even when using a fixed set of gap and substitution scores; different variant calling methods may produce different variant representations. While some of these differences can be handled using pre-processing of VCF files (*e.g.* variant trimming and left-shifting), others cannot be fixed easily. As a result we cannot compare VCF files accurately by comparing VCF records and genotypes directly. Approaches were developed to standardize indel representation by means of left-shifting and trimming the indel alleles.^19,20^ These methods determine the left-most and right-most positions at which a particular indel could be represented in a VCF file (Fig. 2a). These methods work well when considering each VCF record independently. However, when multiple VCF records are used to represent a complex haplotype, normalization methods can cause errors and more sophisticated comparison methods are required (Fig. 2b-d). Different types of variant representation challenges are detailed in Supplementary Information E.

**Fig. 2:**
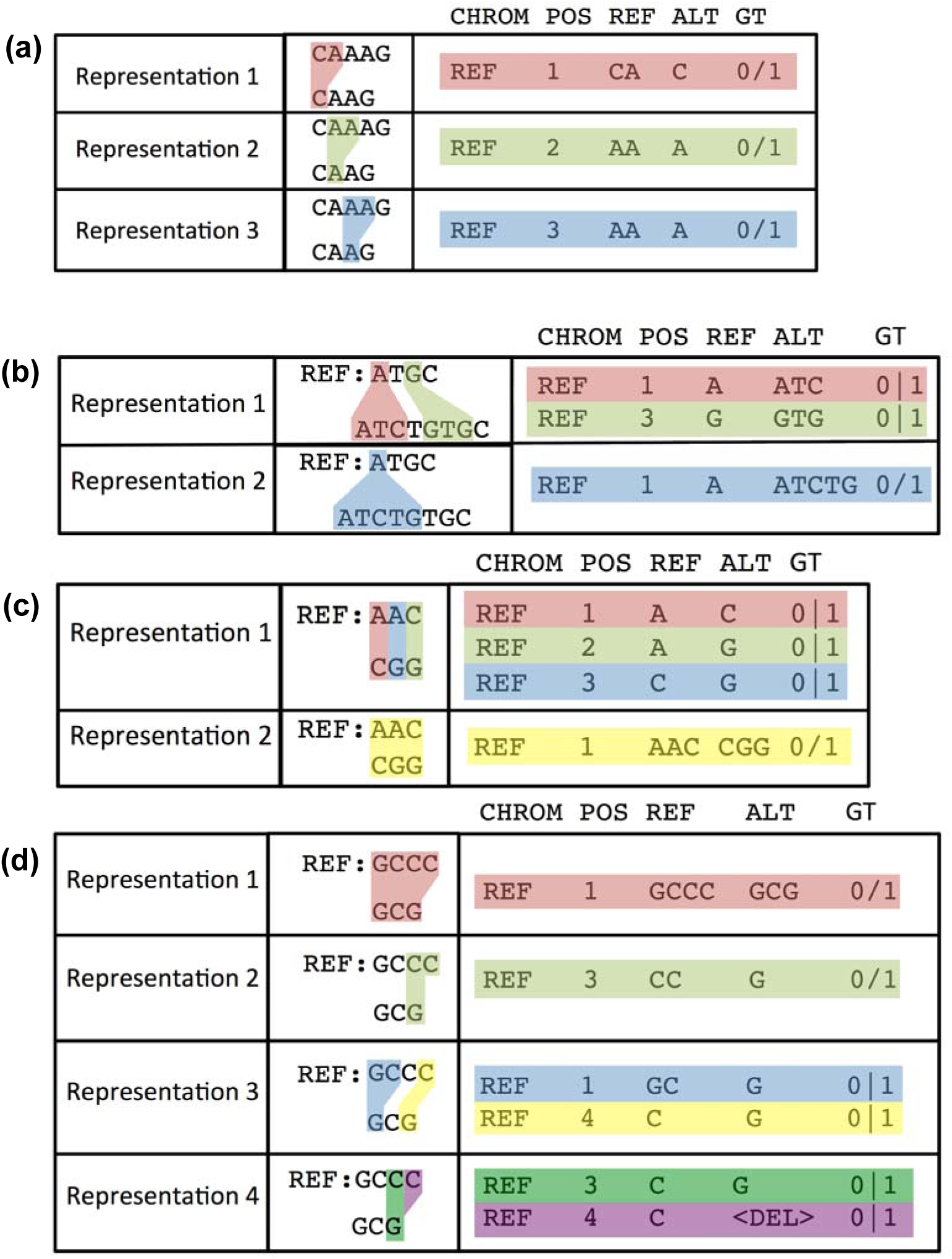
Four examples of cases where variants can be represented in multiple forms in VCF format. (a) Three representations of a deletion in a homopolymer. (b) The insertion can be represented as one 4-bp insertion or two 2-bp insertions. (c) An MNP can be represented as 3 SNVs or one larger substitution. (d) Four different representations of a complex variant. Note that representations include phasing information in these examples where it is necessary to unambiguously describe the variant. If phasing was not described for these variants, it would impossible to normalize their representations, but our sophisticated variant comparison tools can determine that they could describe the same two haplotypes.

When benchmarking, these variant representation differences can also give rise to different notions of giving partial credit for variant calls. One example is where we may have called only one SNV in an MNP with the correct genotype. When assigning TP/FP/FN status on a per-VCF-record basis, a variant caller that chooses to represent calls using single MNP records would not get credit for calling this SNV correctly since the overall MNP record does not reproduce the correct haplotype. Another example would be phasing switch-errors: a choice needs to be made whether to use phasing-aware benchmarking for a particular evaluation. Handling these cases is important since adding phasing information provides additional information to the users of a variant caller, but may lead to FPs / FNs when running a benchmarking comparison when comparing to a method which does not provide phasing information and outputs all alleles in decomposed form for maximum credit in the benchmarking comparison. Our tools attempt to give partial credit when possible, and we generally recommend using vcfeval as the comparator to provide the most partial matches.

## Matching Stringencies and Defining Performance Metrics

Due to the inherent complexity of the human genome, and the challenge that genotype comparisons do not cleanly fall in a binary classification model, TP, FP, and FN can be defined in different ways. Our reference implementation for benchmarking uses a tiered definition of variant matches, a standardized VCF format for outputting matched variant calls, and a common counting and stratification tool (see SI A). We consider three types of variant matches from most to least stringent: (1) “genotype match”, for which only sites with matching alleles and genotypes are counted as TPs, (2) “allele match”, for which any site with matching alleles is counted as TP, even if genotypes differ, and (3) “local match”, for which any site in the query with a nearby truth variant is counted as a TP, even if alleles and genotypes differ. “Genotype match” is used by our current tools to calculate TP, FP, and FN.

In Table 1, we enumerate the types of matches that are clear TP, FP, and FN as well as various kinds of partial matches that may be considered TP, FP, and/or FN depending on the matching stringency, and how they are counted by our tools. Our tools calculate TP, FP, and FN requiring the genotype to match, but output additional statistics related to how many of the FPs and FNs are allele matches (FP.GT) or local matches (FP.AL). Note that we have chosen not to include true negatives (or consequently specificity) in our standardized definitions. This is due to the challenge in defining the number of true negatives, particularly for indels or around complex variants. In addition, precision is often a more useful metric than specificity due to the very large proportion of true negative positions in the genome.

**Table 1:** Contingency table describing the GA4GH definitions of true positive (TP), false positive (FP), false negative (FN), allele mismatch (FP.AL), genotype mismatch (FP.GT), and unknown (UNK). Matches counted as FP.GT and FP.AL are additionally counted as both FP and FN, since our tool’s default matching stringency requires genotypes to match. Query variants outside the Truth bed file are counted as UNK.

|  |  | Truth |  |  |  |  |
| --- | --- | --- | --- | --- | --- | --- |
|  | Genotype | ref/ref | ref/var1 | var1/var2 | var1/var1 | Outside bed |
| Query | ref/ref | - | FN | FN | FN | - |
|  | ref/var1 | FP | TP | <u>FP.GT</u> | <u>FP.GT</u> | UNK |
|  | ref/var2 | - | <u>FP.AL</u> | <u>FP.GT</u> | <u>FP.AL</u> | - |
|  | ref/var3 | - | - | <u>FP.AL</u> | - | - |
|  | var1/var2 | FP | <u>FP.GT</u> | TP | <u>FP.GT</u> | UNK |
|  | var1/var3 | - | - | <u>FP.GT</u> | - | - |
|  | var2/var3 | - | <u>FP.AL</u> | <u>FP.GT</u> | <u>FP.AL</u> | - |
|  | var3/var4 | - | - | <u>FP.AL</u> | - | - |
|  | var1/var1 | FP | <u>FP.GT</u> | <u>FP.GT</u> | TP | UNK |
|  | var2/var2 | - | <u>FP.AL</u> | <u>FP.GT</u> | <u>FP.AL</u> | - |
|  | var3/var3 | - | - | <u>FP.AL</u> | - | - |

To reconcile the comparison methods and metrics discussed above into a simple summary, we have implemented in *hap.py* a standardized report that can be generated from the tabular output of the benchmarking workflow.^21^ An example of the metrics and plots displayed in such a report is shown in Fig. 3. Definitions and formulas for all performance metrics, including derived metrics such as precision and recall, are detailed in the Online Methods and Supplementary Table 1.

**Figure 3:**
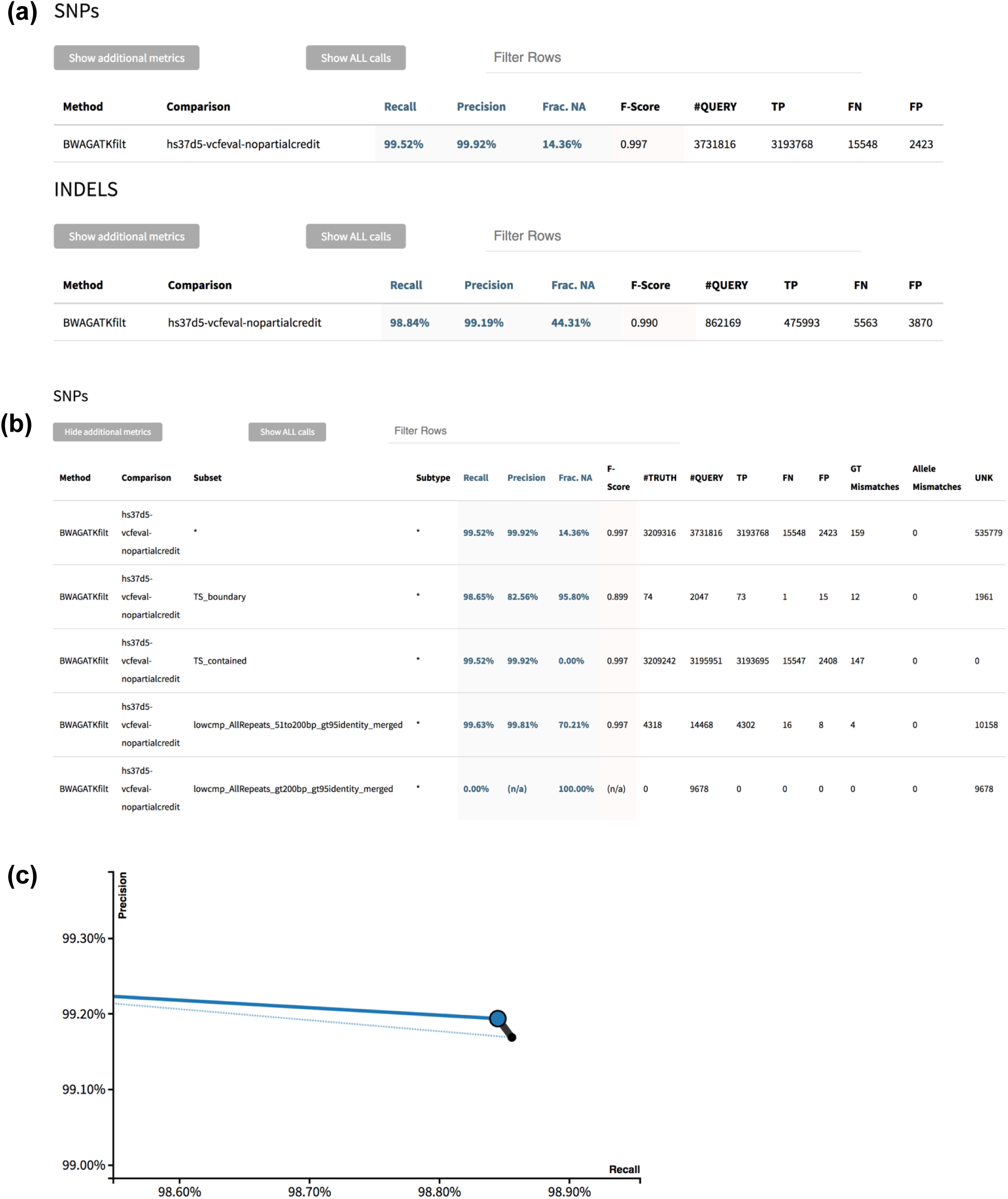
Example standardized HTML report output from hap.py. (a) Tier 1 high-level metrics output in the default view. (b) Tier 2 more detailed metrics and stratifications by variant type and genome context. (c) Precision-recall curve using QUAL field, where the black point is all indels, the blue point is only PASS indels, the dotted blue line is the precision-recall curve for all indels, and the solid blue line is the precision-recall curve for PASS indels.

## Benchmark callsets

Benchmarking of variant calls requires a specific genome and an associated set of calls that represent the “right answers” for that genome. Such call sets have the property that they can be used as “truth” to accurately identify false positives and negatives. That is, when comparing calls from any sequencing method to this set of calls, at least half (and ideally more) of the putative false positives and false negatives should be errors in the method being assessed. Because it is treated as the truth, this benchmark set will be referred to in this manuscript as the “truth” set, but other terms used for this include the “gold-standard” set, the “high-confidence” set, the “reference callset,” or “benchmarking data.”

We describe three sources of benchmark callsets in detail in the Online Methods. Briefly, the Genome in a Bottle Consortium (GIAB) is an ongoing public-private-academic consortium hosted by the National Institute of Standards and Technology (NIST) to perform authoritative characterization of a small number of broadly consented and disseminated human genomes. Currently, five human genomes are available as NIST Reference Materials with benchmark small variant and reference calls for approximately 90% of GRCh37 and GRCh38.^4,6,23^ In addition to the benchmarking data produced by the GIAB consortium, Illumina Platinum Genomes (PG) has also created a benchmarking data set for small variants (SNVs and Indels) using the 17-member pedigree (1463) from Coriell Cell Repositories that includes the GIAB pilot sample NA12878/HG001.^5^ This pedigree includes 11 children of the parents (NA12877 and NA12878), producing a fully phased dataset that allows to validate the accuracy of variant calls through genetic inheritance patterns. Finally, a new “synthetic-diploid” benchmark callset was created from long read assemblies of the CHM1 and CHM13 haploid cell lines, in order to benchmark small variant calls in regions difficult to analyze with short reads or in diploid genomes, which are currently excluded from the GIAB and Platinum Genomes high-confidence regions.^7^ A current limitation is that CHM1 and CHM13 cell lines are not available in a public repository.

## Example comparisons

### Lessons from PrecisionFDA Challenges

The PrecisionFDA team held two challenges in 2016, with participants publicly submitting results from various mapping/variant calling pipelines (more information at https://precision.fda.gov/challenges/). While both challenges asked participants to analyze short read WGS datasets, the first “Consistency” challenge used a sample with high-confidence calls already available (HG001/NA12878) and the second “Truth” challenge used a sample without high-confidence calls yet available (HG002 from GIAB, made available by GIAB upon the close of the challenge).

Note that both the “truth” sets and the comparison methodology in the truth challenge were newly introduced, with GA4GH comparison methodology, truth sets, and variant calling methods under active development. The challenge results available on precisionFDA should be considered only initial evaluation, with the rich data set resulting from the challenge inviting further exploration. It is especially critical to recognize that performance metrics indicate performance for the “easier” variants and regions of the genome, so that precision and recall estimates are higher than if more difficult variants and regions were included. It is likely that some methods will perform worse than other methods for easier variants while performing better for harder variants (*e.g.*, methods using a graph reference or de novo assembly may do better calling in regions not assessed like the MHC or large insertions, while not performing as well for easier variants because the methods are less mature). It is also important to manually curate a subset of FPs and FNs to ensure they are actually FPs and FNs and to understand their cause. Interestingly, stringency of matching can also significantly influence performance metrics. For example, Figure 4 shows how the number of FP indels for the assembly-based fermikit submission is much higher than the RTG submission when counting genotype errors as FPs, but the number of FPs is lower for fermikit when matching only the allele or performing distance-based matching. Additional information about relative strengths and weaknesses of the pipelines could also be gained through stratification, as discussed in the next section.

**Figure 4:**
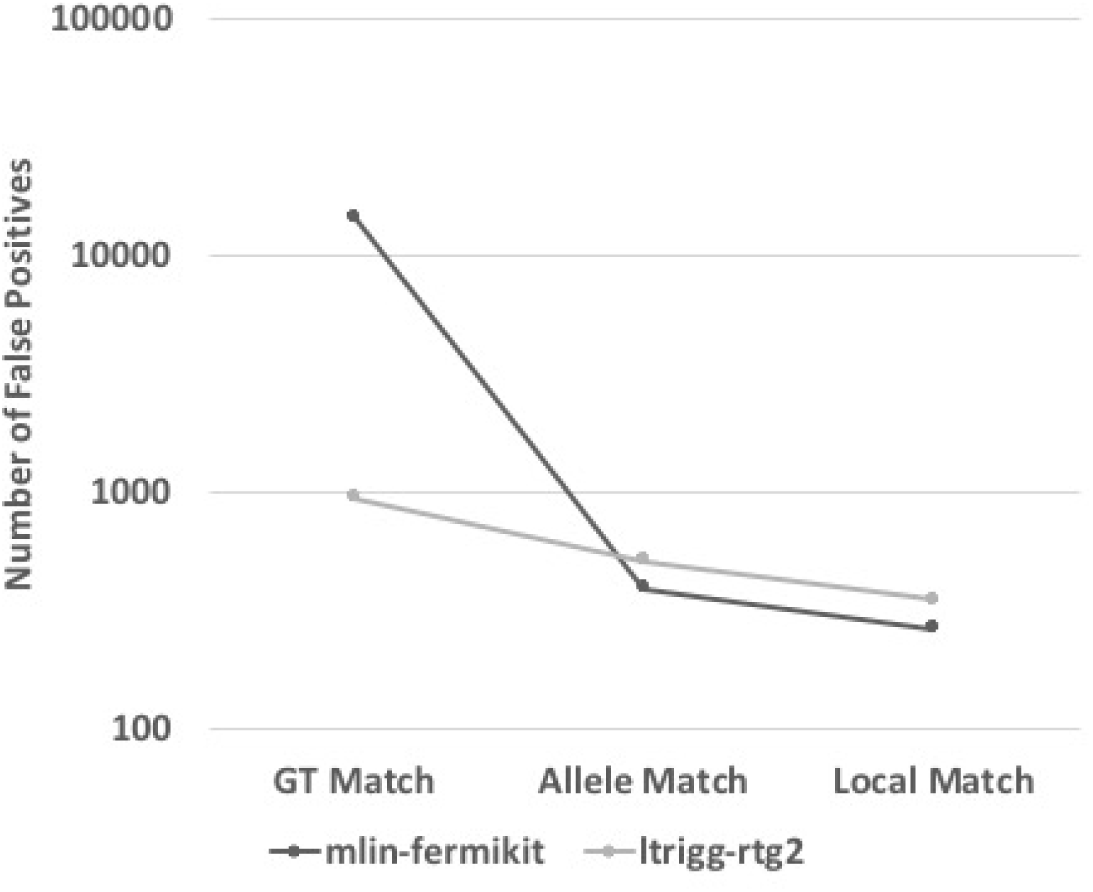
Matching stringency can affect relative performance of algorithms. Number of false positives for two PrecisionFDA Challenge submissions is shown for different matching stringencies, showing that the fermikit submission has many more false positives if genotype errors are counted as FPs, but that it has fewer FPs if matching only the allele or performing distance-based matching. Note that this is intended to illustrate the importance of matching stringency and is likely not indicative of the performance of these methods with optimized parameters or current versions.

## Stratification illuminates challenging regions sequenced with and without PCR amplification

Our team has defined a large number of regions of different genome contexts (e.g., GC content and repeats of different sizes and types) to enable users to stratify performance and understand strengths and weaknesses of a particular method. As an example of using stratification, we compare recall and precision in different genome contexts for whole genome sequencing assays with and without a PCR amplification step. Table 2 shows that indel recall and precision are lower when using PCR amplification than when using PCR-free sequencing. Stratification highlights that this difference almost entirely results from PCR-related errors in homopolymers and tandem repeats, since performance is similar when excluding variants that occur within 5bp of homopolymer sequences longer than 5bp and tandem repeats longer than 10bp. Performance in regions with low GC content is similar, but PCR results in lower SNV and indel recall where GC content is > 85%.

**Table 2:**
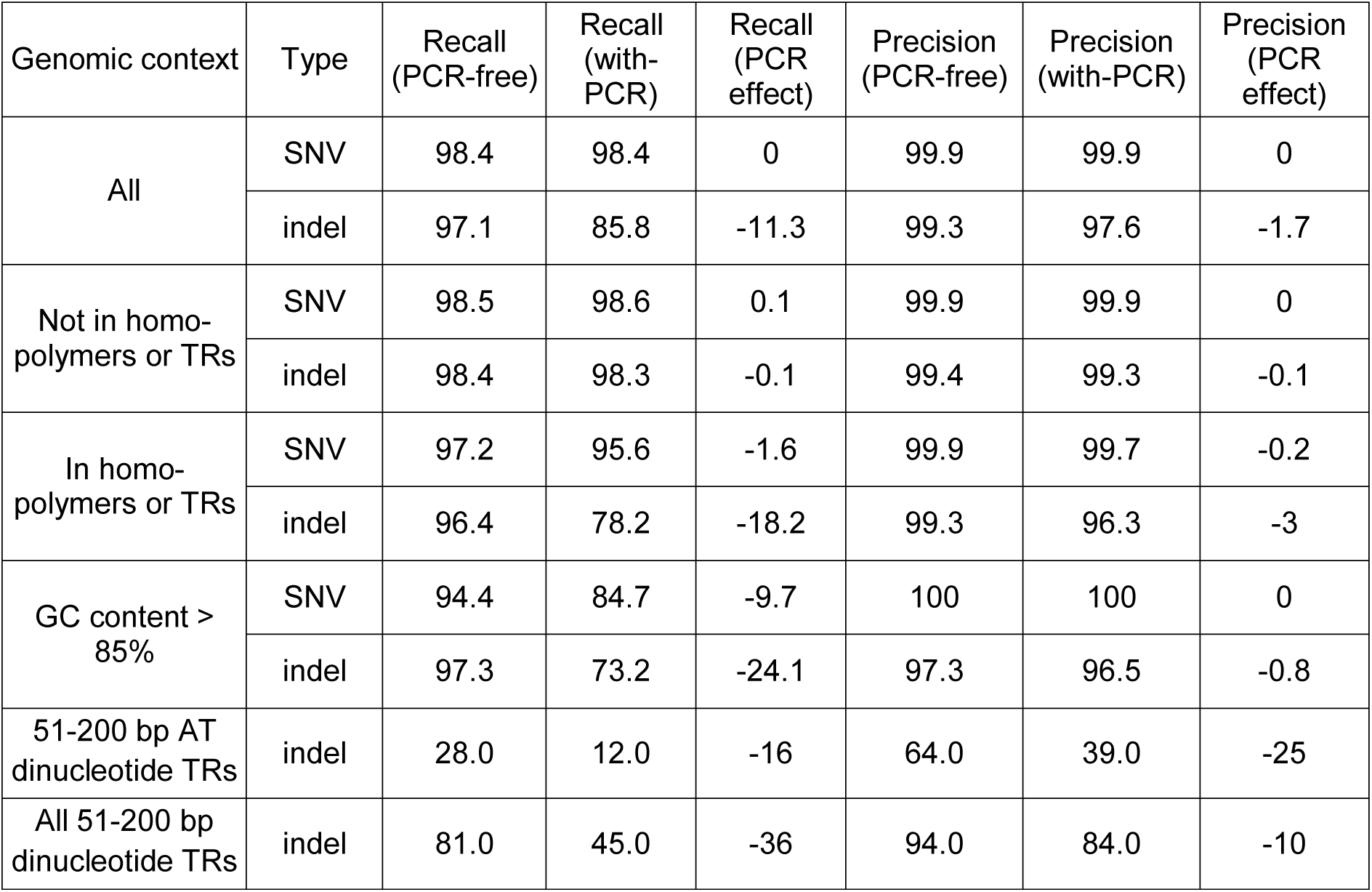
Recall and Precision stratified by genomic context (e.g., GC content and tandem repeat (TR) type) and variant type for Illumina whole genome sequence assays with and without a PCR step

Further stratification by type of repeat can illuminate particularly challenging genome contexts. For example, when sorting strata by recall, indels in 51-200 bp AT dinucleotide tandem repeats have substantially lower recall and precision than all other strata for both PCR and PCR-free results. Also, 86 out of 114 truth indels in 51-200 bp AT dinucleotide tandem repeats are compound heterozygous, and 89% fall outside the high-confidence regions, so our stratification and benchmarking methods help illuminate that these appear to be highly polymorphic and difficult variants to characterize.

## Benchmarking Best Practices

Box 1: GA4GH recommendations for best practices for germline variant call benchmarking

|  |  |
| --- | --- |
| **Benchmark sets** | Use benchmark sets with both high-confidence variant calls as well as high-confidence regions (e.g., from GIAB or Platinum Genomes). |
| **Stringency of variant comparison** | Determine whether it is important that the genotypes match exactly, only the alleles match, or the call just needs to be near the true variant. For example, if you confirm and/or manually curate all variants to ensure you have the correct allele and genotype, then local matching may be sufficient. While the default TP, FP and FN require genotype and allele matching, the additional metrics FP.GT and FP.AL output by the GA4GH tools enable users to calculate performance at different stringencies. |
| **Variant comparison tools** | Use sophisticated variant comparison engines such as vcfeval, xcmp, or varmatch that are able to determine if different representations of the same variant are consistent with the benchmark call (examples in Fig. 1). Subsetting by high-confidence regions and, if desired, targeted regions, should only be done after comparison to avoid problems comparing variants with different representations. |
| **Manual curation** | Manually curate alignments, ideally from multiple data types, around at least a subset of putative false positive and false negative calls in order to ensure they are truly errors in the user's callset and to understand the cause(s) of errors. Report back to benchmark set developers any potential errors found in the benchmark set (e.g., using https://goo.gl/forms/ECbjHY7nhz0hrCR52 for GIAB or https://github.com/Illumina/PlatinumGenomes/issues/new for PG). |
| **Interpretation of metrics** | All performance metrics should only be interpreted with respect to the limitations of the variants and regions in the benchmark set. Performance is unknown for variant types and genome contexts not well represented in the benchmark set. Performance metrics are likely to be lower for more difficult variant types and regions that are not fully represented in the benchmark set, such as those in repetitive or difficult-to-map regions. When comparing methods, note that method A may perform better in the high-confidence regions, but method B may perform better for more difficult variants outside the high-confidence regions. |
| **Stratification** | Performance results should be stratified by variant type. Stratification by genomic region should also be considered to gain additional insights into strengths and limitations of the sequencing pipeline, as it can highlight regions that are not sufficiently represented. Stratification should only be done after comparison to avoid problems comparing variants with different representations. |
| **Confidence Intervals** | Confidence intervals for performance metrics such as precision and recall should be calculated. This is particularly critical for the smaller numbers of variants found when benchmarking targeted assays and/or less common stratified variant types and regions. |
| **Additional benchmarking approaches** | We recommend using other benchmarking approaches in addition to those discussed in this paper to understand performance of a pipeline, including:<br>• Confirming results found in samples over time<br>• Synthetic DNA spike-ins with challenging and common clinically relevant variants<br>• Engineering variants into cell lines<br>• Finding existing samples with challenging and common clinically relevant variants<br>• Simulation methods, such as read simulators, adding variants into real reads, and modifying the reference<br>• Run-specific metrics such as base quality score distributions, coverage distributions, etc. can also be useful to identify outlier runs |

## Conclusions

The GA4GH Benchmarking Team has developed a suite of methods to produce standardized performance metrics for benchmarking small germline variant calls. These sophisticated tools address challenges in standardizing metrics like recall and precision, comparing different representations of variant calls, and stratifying performance by variant type and genome context. We have developed a set of best practices for benchmarking variant calls to help users avoid common pitfalls and misinterpretations of performance metrics.

Moving forward there will be a continual need for improvements in benchmarking of variant discovery methodologies. Technological evolution will enable laboratories to characterize increasingly difficult variants and genomic regions, which will require improved benchmarks. Simultaneously, this evolution can contribute to improved characterization of reference materials through ongoing work by groups like GIAB. For example, the types of variants being analyzed will increase in scope: most current benchmarking focuses on relatively small variations, and quite different techniques will be needed to consider structural variants. In addition to the genotype, allele, and local matching stringencies we describe for small variants, comparison tools for structural variants will need to consider stringencies for breakpoint matching, size predictions, and inserted sequence predictions.

Assessment of somatic variants also introduces challenges different from germline variants. Different benchmarking approaches are needed to handle somatic issues like assessing the accuracy of variant allele frequency. A good germline variant caller will not perform well for somatic detection, and *vice versa*. A global consortium around benchmarking of somatic variant detection has been established, called the ICGC-TCGA DREAM Somatic Mutation Calling (SMC) group, and has been benchmarking both detection of individual variants and of broader processes like subclonal variation.^24^

Moving forward, groups will also need to modify benchmarking strategies to address changes in the way the human genome itself is represented. Today the most common way of representing the human genome involves a set of linear chromosomes (*e.g.,* the most common usage of GRCh37). There are key advantages to non-linear representations of the genome, including ability to recognize copy-number and other polymorphisms directly in the reference, and as a result more graphical structures are in development.^25^ The GRCh38 build of the human genome makes a key step towards this with its use of ALT loci, which provide multiple distinct versions of specific regions of the genome.^26^ These ALT loci are not well-accounted for by most aligners or the benchmarking tools we describe, and their impact on benchmarking studies is largely unexplored and likely would require a variety of samples with differing ALT alleles. It is likely that the core representation of the genome will continue to evolve over time, and benchmarking tools will continue to evolve.

This work provides a framework of principles for further development of benchmarking tools to address new challenges in variant calling and other high-throughput measurement challenges.

## Acknowledgments

We thank GA4GH, especially Stephen Keenan, David Lloyd, and Rishi Nag, for their support in hosting and organizing the Benchmarking Team. We thank the many contributors to Benchmarking Team and Genome in a Bottle Consortium discussions over the past few years, especially Deanna Church, Kevin Jacobs, and Brendan O’Fallon. Certain commercial equipment, instruments, or materials are identified to specify adequately experimental conditions or reported results. Such identification does not imply recommendation or endorsement by the National Institute of Standards and Technology or the Food and Drug Administration, nor does it imply that the equipment, instruments, or materials identified are necessarily the best available for the purpose.

## Author contributions

PK, LT, PCB, CEM, FMD, MAE, RT, BF, MF, MS, and JMZ wrote the manuscript.

PK, LT, FMD, BLM, and MG designed and implemented the benchmarking tools.

ZT, SL, GA, and JMZ designed and/or analyzed results from PrecisionFDA Challenges

PK, LT, GA, BC, MS, and JMZ designed the project

All authors contributed to GA4GH Benchmarking Team discussions about this work.

## Online Methods

### Variant representation

A variety of approaches have been recently developed to address the challenges in variant representation.^9–11,21,22^ Real Time Genomics (RTG) developed the comparison tool *vcfeval*, which introduced the idea of comparing variants at the level of the genomic haplotypes that the variants represent as a way to overcome the problems associated with comparing complex variants, where alternative yet equivalent variant representations can confound direct comparison methods.^9^ Variant “normalization” tools help to represent variants in a standardized way (e.g., by left-shifting indels in repeats), but they demonstrated that “variant normalization” approaches alone were not able to reconcile different representations of many complex variants. In contrast, global optimization permits evaluation of alternate representations that minimize the number of discrepancies between truth and test set caused by differences in representations of the same variant. Similarly, *VarMatch* was developed to resolve alternate representations of complex variants, with additional ability to tune the matching parameters depending on the application.^10^ Finally, *hap.py* includes a comparison tool to perform haplotype-based comparison of complex variants in addition to sophisticated functionality to stratify variant calls by type or region.^21^ We use the *hap.py* framework with the *vcfeval* comparison tool in this work.

### Variant counting

The GA4GH Benchmarking Team developed consensus definitions and recommendations for expressing performance metrics for small germline variant calls. Assessing the performance of variant callers does not easily lend itself to the typical binary classification performance assessment model of simply determining true and false “positives” and “negatives”. Several characteristics of the genome do not fit well in a binary classification model:

1. More than two possible genotypes exist at any given location. For SNVs (if ignoring phasing), any location can have one of 10 different true genotypes (*i.e.*, A/A, A/C, A/G, A/T, C/C, C/G, …). For indels and complex variants, an infinite number of possible genotypes exists (*e.g.*, any length of insertion).
2. A number of variant callers distinguish between “no-calls” and homozygous reference calls at some genome positions or regions. Some variant callers even output partial no-calls, calling one allele but not the other. “No-calls” at a true variant site could be treated as false negatives or be excluded from counting.
3. In addition to the challenges comparing different representations of complex variants (*i.e.*, nearby SNVs and/or indels) discussed above, there are challenges in standardizing counting of these variants. Complex variants can be treated as a single positive event or as multiple distinct SNV and indel events when counting the number of TP, FP, and FN variants. In addition, only part of a complex variant may be called, which poses challenges in defining TP, FP, and FN.
4. Methods for assessing accuracy of phasing have not been fully developed or standardized, but accurate phasing can be critical, particularly when multiple heterozygous variants exist in a small region (*e.g.*, complex variants).

### Matching Stringencies

Due to the inherent complexity of the human genome, TP, FP, and FN can be defined in different ways. Our reference implementation for benchmarking uses a tiered definition of variant matches, a standardized VCF format for outputting matched variant calls, and a common counting and stratification tool (see SI A).

We consider the following types of variant matches from most to least stringent, with “Genotype match” being used by our current tools to calculate TP, FP, and FN:

- *Genotype match*: Variant sets in truth and query are considered TPs when their unphased genotypes and alleles can be phased to produce a matching pair of haplotype sequences for a diploid genome. Each truth (and query) variant may be replayed onto one of two truth (or query) haplotypes. A maximal subset of variants that is replayed to produce matching haplotype sequences forms the TP variants, query variants outside this set are FP, truth variants outside this set are FN. The method only considers haploid or diploid samples but could be extended to higher ploidy also. Enumerating the possible assignments for haplotype generation is computationally expensive. Vcfeval solves this problem using global optimization methods supplemented with heuristic pruning. Genotype match statistics are the default TP, FP, and FN output by our tools. Genotype matching has been implemented in the *hap.py* tool *xcmp* and in *vcfeval*.
- *Allele match*: Truth and query alleles are counted as TP_AM if they contain any of the same (trimmed and left-shifted) alleles. This method is more specific than local matching (e.g. repeat expansions must be called with the correct length in order to get an allele match), but could also be susceptible to spurious mismatches when truth and query variant alleles are decomposed differently. Genotype mismatches (FP.GT in Table 1) are considered TPs in this matching method. We indicate allele matches in scenarios where variants can be matched when ignoring the genotype. Allele match statistics (TP_AM, FP_AM, and FN_AM) can be calculated from the GA4GH outputs (which require genotypes to match): TP_AM=QUERY.TP+FP.GT; FP_AM=QUERY.FP-FP.GT; FN_AM=TRUTH.FN-FP.GT. Allele matching has been implemented in the *hap.py* tool *scmp-somatic* and in *vcfeval* with the --squash-ploidy option.

- Note that *vcfeval* --squash-ploidy and *scmp-somatic* differ. *scmp-somatic* checks if the VCF records give the same alleles after normalization and trimming. This will match alleles that overlap on the reference as long as they can be matched directly after left-shifting and trimming. When comparing somatic variant calls, this is probably the best option since technically, every variant could be on a different (low-frequency) haplotype. *vcfeval* --squash-ploidy does haplotype-based comparison but assumes all variants are hom-alt and there is only one haplotype. This will match different representations unless they overlap on the reference (which is also possible using *xcmp* via the force-gt command line option in *hap.py* which changes the GTs before comparing).
- *Local match*: Truth and query variants are counted as TP_LM if their reference span intervals are closer than a pre-defined local matching distance, i.e. all yellow categories in Table 1 are considered TPs, including “F” matches that are within a specified number of basepairs. This approach has previously been implemented.^7,21^ An advantage of this matching method is that it is robust towards representational differences. A drawback for many applications is that it does not measure allele or genotype accuracy. We use local matches as the lowest tier of matching to label variants which are close-by but cannot be matched with other methods. Local match statistics (TP_LM, FP_LM, and FN_LM) can be calculated from the GA4GH outputs (which require genotypes to match): TP_LM=QUERY.TP+FP.GT+FP.AL; FP_LM=QUERY.FP-FP.GT-FP.AL; FN_LM=TRUTH.FN-FP.GT-FP.AL. If only local matching is required, this has been implemented in the *hap.py* tool *scmp-distancebased.*

A fourth, most stringent matching, which is not yet fully implemented in the GA4GH framework, requires phasing information to match:

- *Phased genotype match*: When VCF files specify phasing information, we can compare on a haplotype level: variants will only be matched if they produce matching haplotype sequences under phasing constraints. Both vcfeval and hap.py’s xcmp method support phased matching when both callsets include variants that are globally phased (*i.e.* specify a paternal and maternal haplotype for each chromosome). To our knowledge, no current comparison method supports phasesets and local phasing to compare variants. Moreover, assessing phasing requires us to consider not only phasing variant accuracy, but also completeness of phasing coverage. In our current methods we do not implement phased genotype matching beyond the basic support provided by vcfeval and xcmp.

### Defining True Positives, False Positives, and False Negatives

In Table 1, we enumerate the types of matches that are clear TP, FP, and FN as well as various kinds of partial matches that may be considered TP, FP, and/or FN depending on the matching stringency, and how they are counted by our tools. Our tools calculate TP, FP, and FN requiring the genotype to match, but output additional statistics related to how many of the FPs and FNs are allele matches (FP.GT) or local matches (FP.AL). Note that we have chosen not to include true negatives (or consequently specificity) in our standardized definitions. This is due to the challenge in defining the number of true negatives, particularly around complex variants. In addition, precision is often a more useful metric than specificity due to the very large proportion of true negative positions in the genome.

Another key question is how to count both matching and mismatching variant calls when they are differently represented in the truth dataset and a query. When representing MNPs as multiple SNVs, we may count one variant call for each SNV, or only one call in total for the MNP record. Similar considerations apply to counting complex records. We approach variant counting as follows:

- We count the truth and query VCF files separately. A set of truth records may be represented by a different set of query records.
- To get comparable recall, we count both TPs and FNs in their truth representation. When comparing different variant calling results to the same truthset, these counts will be based on the same variant representation.
- Precision is assessed using the query representation of variants. We give a relative precision to the number of truth variants in query representation. If a variant caller is consistent about the way it represents variants, this approach mitigates counting-related performance differences.
- We implement a “partial credit” mode in which we trim, left-shift and decompose all query variant calls before comparison. This resolves the MNP vs. SNV comparison issues and also simplifies the variant types we use for stratification, rather than having a category of complex variant calls which has results that are difficult to interpret, we account for every atomic indel and SNV call independently.
- Variants are stratified into a canonical set of types and subtypes (see SI B).
- When stratification regions are applied, we match variants by their trimmed reference span. If any part of a deletion overlaps the stratification region, it is counted as part of that stratum. Insertions receive special treatment by requiring both the base before and the base after to be captured. Importantly, this stratification is performed after comparison to deal appropriately with representation issues.

### Benchmarking metrics report

To reconcile the comparison methods and metrics discussed above into a simple summary, we have implemented in *hap.py* a standardized report that can be generated from the tabular output of the benchmarking workflow.^21^ This report displays the metrics we believe are most important in an accessible fashion (Tier 1 metrics), while also allowing to examine the data in more detail (Tier 2 metrics). An example for the metrics and plots displayed in such a report is shown in Fig. 3.

From the TP, FP, and FN counts defined in Table 1, we calculate:

METRIC.PRECISION = QUERY.TP / (QUERY.TP + QUERY.FP)

METRIC.RECALL = TRUTH.TP / (TRUTH.TP + TRUTH.FN)

We use the count of TPs based on the query representation (QUERY.TP) to calculate precision, and we use the count of TPs based on the Truth representation (TRUTH.TP) to calculate recall, in order to account best for cases where the Truth may tend to split a complex variant into multiple varaints and the Query may combine them into a single variant, or vice versa. Definitions and formulas for all performance metrics are detailed in Supplementary Table 1.

An alternative to precision is false positive rate (FPR) = FP / megabase.^7^ It can easily be obtained from GA4GH/hap.py extended csv by taking FP / 1e6 * Subset.Size (or Subset.IS_CONF.Size, the number of confident bases in each stratification region). Precision approximates the probability that a given query call is true, while FPR approximates the probability of making a spurious call. Note that we do not define “True negatives” or “specificity” because these are not cleanly applicable to genome sequencing. For example, there are an infinite number of possible indels in the genome, so there are an infinite number of true negatives for any assay.

In addition, the GA4GH Benchmarking framework is able to produce precision-recall curves, which are graphical plots that illustrate the performance of a variant quality score of a test call set as its discrimination threshold is varied, compared to the reference call set (see Figure 4). The curve is created by plotting the precision against the recall at various quality score threshold settings. Commonly used quality scores include QUAL, GQ (genotype quality), DP (depth of coverage), and machine-learning derived scores such as VQSLOD and AVR. Because some methods use multiple annotations for filtering, precision-recall curves can be generated for a particular quality score either before or after removing filtered sites. Examining the precision-recall curves for various call-sets has two main advantages. Firstly, it allows the user to consider how accuracy is affected through the precision/recall trade-off. Secondly, different call sets may have effectively selected different precision/recall trade-off criteria, so simply comparing full call set metrics may reflect more about the different trade-off points than the call sets themselves at some shared trade-off criteria.

### Benchmark callsets

Benchmarking of variant calls requires a specific genome and an associated set of calls that represent the “right answers” for that genome. Such call sets have the property that they can be used as “truth” to accurately identify false positives and negatives. That is, when comparing calls from any sequencing method to this set of calls, >50% of the putative false positives and false negatives should be errors in the method being assessed. Because it is treated as the truth, this benchmark set will be referred to in this manuscript as the “truth” set, but other terms used for this include the “gold-standard” set, the “high-confidence” set, the “reference callset,” or “benchmarking data.”

#### Genome in a Bottle

The Genome in a Bottle Consortium (GIAB) is a public-private-academic consortium hosted by the National Institute of Standards and Technology (NIST) to perform authoritative characterization of a small number of human genomes to be used as benchmarks. GIAB published a benchmark set of small variant and reference calls for its pilot genome, NA12878, which characterized a high-confidence genotype for approximately 78% of the bases with sequence information (i.e., bases that are not an “N”) in the human genome reference sequence (version GRCh37).^3^ Since this publication, GIAB has further developed integration methods to be more reproducible, comprehensive, and accurate, and has incorporated new technologies and analysis methods. The new integration process has been used to form benchmark small variant and reference calls for approximately 90% of GRCh37 and GRCh38 for NA12878, as well as a mother-father-son trio of Ashkenazi Jewish ancestry and the son in a trio of Chinese ancestry from the Personal Genome Project (v3.3.2 at ftp://ftp-trace.ncbi.nlm.nih.gov/giab/ftp/release/).^4,6,23^ The five GIAB-characterized genomes are available as NIST Reference Materials (RMs 8391, 8392, 8393, and 8398), which are extracted DNA from a single, homogenized, large growth of cells for each genome. These samples are also all available as cell lines and DNA from the Coriell Institute for Medical Research. The Personal Genome Project samples are also consented for commercial redistribution,^23^ and several derived products are commercially available, including FFPE-preserved and *in vitro* mutated cell lines, or with DNA spike-ins with particular variants of clinical interest. GIAB is continuing to improve the characterization of these genomes to characterize increasingly difficult variants and regions with high-confidence.

#### Platinum Genomes

In addition to the benchmarking data produced by the GIAB consortium, Illumina Platinum Genomes (PG) has also created a benchmarking data set for small variants (SNVs and Indels) using the 17-member pedigree (1463) from Coriell Cell Repositories that includes the GIAB pilot sample NA12878/HG001.^5^ Every sample of this pedigree was sequenced to ∼50x depth on an Illumina HiSeq2000 system. Variant calls were made from this data using different combinations of aligners and variant callers. This pedigree includes 11 children of the parents (NA12877 and NA12878), producing a fully phased dataset that allows to validate the accuracy of variant calls through genetic inheritance patterns. The HiSeq2000 sequence data used to create these benchmarking calls can be obtained from the Database of Genotypes and Phenotypes (dbGaP; https://www.ncbi.nlm.nih.gov/gap) under accession number phs001224.v1.p1. Additionally, the sequence data for six of the members of this pedigree are released through the European Nucleotide Archive (ENA; http://www.ebi.ac.uk/ena) under accession number ERP001960. The DNA and cell lines for all samples are available from the Coriell Institute for Medical Research, and DNA from a single, homogeneous batch of NA12878 is also available as NIST Reference Material 8398.

#### Merged PG and GiaB

Since the two resources mentioned above constitute two different methods for generating “truth” call sets for NA12878, we have merged these into a single and more comprehensive dataset. Such a “hybrid” truth set can leverage the strengths of each input, namely the diversity of technologies used as input to Genome in a Bottle and the robust validation-by-inheritance methodology employed by Platinum Genomes.

As a first pass, we have compared the call sets in NA12878 and identified the intersection as well as the ones unique to each (Supplementary Figure 2). Next, starting from the union, we have used a modified version of the k-mer validation algorithm described in [PG] to validate the merged calls (Supplementary Methods). This hybrid benchmark call set includes more total variants than either input set (67-333k additional SNVs and 85-90k additional indels), allowing us to assess more of the calls made by any sequencing pipeline without a loss in precision (see below).

This new benchmarking set represents the first step towards a more comprehensive call set that includes both “easy” to characterise variants and those that occur in difficult parts of the genome. Despite this significant advance, there remain areas for continued improvement, such as adjudication between conflicting calls and the merging of confident regions. We will continue to develop this integration method in order to further expand the breadth of coverage of this hybrid truth set resource.

Currently, neither PG nor GIAB makes high-confidence calls on chromosome Y or the mitochondrial genome. In addition, GIAB currently has chromosome X calls only for females, but PG has haploid chromosome X calls for the male NA12877 as well. Hap.py has an optional preprocessing step to guess male/female from the truth VCF. For male samples it converts haploid 1 GT calls on chrX/Y to 1/1 so that they get compared correctly by xcmp. For vcfeval, haploid 1 GT calls are treated as the same as 1/1, so this conversion is not necessary.

#### Synthetic Diploid

A new “synthetic-diploid” benchmark callset was created from long read assemblies of the CHM1 and CHM13 haploid cell lines, in order to benchmark small variant calls in regions difficult to analyze with short reads or in diploid genomes, which are currently excluded from the GIAB and Platinum Genomes high-confidence regions.^7^ Because it is based on long reads, performance metrics are likely less biased toward any short read sequencing technology or informatics method, and it enables benchmarking in regions difficult to map with short reads. However, because it currently contains some errors that were not corrected in the long reads, it requires a less stringent benchmarking methodology similar to the “local match” method described below. It also excludes 1bp indels from performance assessment since long read assemblies contain 1bp indel errors, and >50bp indels because these are not analyzed. Therefore, it is currently not as useful for assessing accuracy of genotypes or accuracy of the exact sequence change predicted in the REF and ALT fields. When using GA4GH tools requiring genotypes to match, the majority of FPs and FNs may not be errors in the query callset, though work is underway to improve this. Nevertheless, it is likely to be complementary to the GA4GH benchmarking strategy by enabling users to assess accuracy in more difficult regions that GIAB and Platinum Genomes currently exclude from their high confidence regions. In particular, because the truth set was not developed from short reads, and errors in the truth may be different from errors in short reads, it may better assess of relative performance between short read-based methods, particularly in more difficult genomic regions. A current limitation is that CHM1 and CHM13 cell lines are not available in a public repository.

### PrecisionFDA Challenges

The PrecisionFDA team held two challenges in 2016, with participants publicly submitting results from various mapping/variant calling pipelines. While both challenges asked participants to analyze short read WGS datasets, the first challenge used a sample with high-confidence calls already available (HG001/NA12878) and the second one without high-confidence calls yet available (HG002 from GIAB, made available by GIAB upon the close of the challenge).

In the first, “Consistency” Challenge, 30x Illumina WGS of the HG001/NA12878 sample was provided from two different sequencing sites, and the VCF file results from 17 participants were assessed for reproducibility and accuracy against the GIAB v2.19 Benchmark VCF. It is possible to generate reproducible results without much variability but substantial differences from the truth. Additionally, the pipelines that generated the variant calls could be tuned to HG001, which, in many situations, was used to train or optimize pipelines.

Therefore, in the second, “Truth” Challenge, participants were asked to use their pipelines with 50x Illumina WGS to predict variants from at the time yet unknown reference sample HG0002/NA24385. Challenge results were compared using two benchmarking comparator tools, RTG Tools vcfeval for Consistency Challenge, and Vcfeval + Hap.py Comparison for Truth Challenge (more information at https://precision.fda.gov/challenges/). There were 35 entries in the Truth Challenge and the responses were submitted and ranked according to precision and recall for SNVs and indels vs. the GIAB v3.3.2 high-confidence calls for each genome (ftp://ftp-trace.ncbi.nlm.nih.gov/giab/ftp/release/). This was the first time Vcfeval + Hap.py GA4GH comparison methodology was applied at scale across the large number of entries submitted by pipeline developers. It helped highlight the utility of the tools and the need for further development and careful interpretation of results. Based in part on feedback from the challenges, an improved benchmarking app “GA4GH Benchmarking” uploaded by user peter.krusche is now available on precisionFDA.

## References

1. Yang, Y. et al. Molecular Findings Among Patients Referred for Clinical Whole-Exome Sequencing. JAMA 312, 1870 (2014).

2. Xue, Y., Ankala, A., Wilcox, W. R. & Hegde, M. R. Solving the molecular diagnostic testing conundrum for Mendelian disorders in the era of next-generation sequencing: single-gene, gene panel, or exome/genome sequencing. Genet. Med. 17, 444–451 (2015).

3. Zook, J. M. et al. Integrating human sequence data sets provides a resource of benchmark SNP and indel genotype calls. Nat. Biotechnol. 32, 246–51 (2014).

4. Zook, J. M. et al. Extensive sequencing of seven human genomes to characterize benchmark reference materials. Sci. Data 3, 160025 (2016).

5. Eberle, M. A. et al. A reference data set of 5.4 million phased human variants validated by genetic inheritance from sequencing a three-generation 17-member pedigree. Genome Res. (2016). DOI:10.1101/gr.210500.116

6. Zook, J. et al. Reproducible integration of multiple sequencing datasets to form high-confidence SNP, indel, and reference calls for five human genome reference materials. bioRxiv 281006 (2018). DOI:10.1101/281006

7. Li, H. et al. New synthetic-diploid benchmark for accurate variant calling evaluation. bioRxiv 223297 (2017). DOI:10.1101/223297

8. Highnam, G. et al. An analytical framework for optimizing variant discovery from personal genomes. Nat. Commun. 6, 6275 (2015).

9. Cleary, J. G. et al. Comparing Variant Call Files for Performance Benchmarking of Next-Generation Sequencing Variant Calling Pipelines. bioRxiv (Cold Spring Harbor Labs Journals, 2015). DOI:10.1101/023754

10. Sun, C. & Medvedev, P. VarMatch: robust matching of small variant datasets using flexible scoring schemes. Bioinformatics 33, btw797 (2016).

11. Talwalkar, A. et al. SMaSH: A benchmarking toolkit for human genome variant calling. Bioinformatics 30, 2787–2795 (2014).

12. The Variant Call Format Specification. (2017). at <https://samtools.github.io/hts-specs/VCFv4.3.pdf>

13. Roper, W. L. et al. Good Laboratory Practices for Molecular Genetic Testing for Heritable Diseases and Conditions. MMWR 58, (2009).

14. Mattocks, C. J. et al. A standardized framework for the validation and verification of clinical molecular genetic tests. Eur. J. Hum. Genet. 18, 1276–1288 (2010).

15. Gargis, A. S. et al. Assuring the quality of next-generation sequencing in clinical laboratory practice. Nat. Biotechnol. 30, 1033–1036 (2012).

16. Rehm, H. L. et al. ACMG clinical laboratory standards for next-generation sequencing. Genet. Med. 15, 733–747 (2013).

17. Aziz, N. et al. College of American Pathologists’ Laboratory Standards for Next-Generation Sequencing Clinical Tests. Arch. Pathol. Lab. Med. 139, 481–493 (2015).

18. Roy, S. et al. Standards and Guidelines for Validating Next-Generation Sequencing Bioinformatics Pipelines: A Joint Recommendation of the Association for Molecular Pathology and the College of American Pathologists. J. Mol. Diagn. 20, 4–27 (2018).

19. Hasan, M. S., Wu, X., Watson, L. T., Li, Z. & Zhang, L. UPS-indel: A Universal Positioning System For Indels. bioRxiv (2017). at <http://biorxiv.org/content/early/2017/05/03/133553>

20. Tan, A., Abecasis, G. R. & Kang, H. M. Unified representation of genetic variants. Bioinformatics 31, 2202–2204 (2015).

21. Krusche, P. Haplotype comparison tools / hap.py. at <http://github.com/illumina/happy>

22. Jacobs, K. B. Variant Graph Comparison Tool (vgraph). at <https://github.com/bioinformed/vgraph>

23. Ball, M. P. et al. A public resource facilitating clinical use of genomes. Proc. Natl. Acad. Sci. U. S. A. 109, 11920–7 (2012).

24. Ewing, A. D. et al. Combining tumor genome simulation with crowdsourcing to benchmark somatic single-nucleotide-variant detection. Nat. Methods 12, 623–30 (2015).

25. Novak, A. M. et al. Genome Graphs. bioRxiv 101378 (2017). DOI:10.1101/101378

26. Schneider, V. A. et al. Evaluation of GRCh38 and de novo haploid genome assemblies demonstrates the enduring quality of the reference assembly. Genome Res. 27, 849–864 (2017).

